# Development of antibacterial drug plus bacteriophage combination assays

**DOI:** 10.1101/2024.03.07.583836

**Authors:** Marie Attwood, Pippa Griffin, Alan Noel, Theo Josephs, Karen Adler, Martha Clokie, Alasdair MacGowan

**Author notes:** Corresponding author Marie Attwood.

## Abstract

**Background:** The methods to evaluate the interactions between Phages and antibacterials are unclear. As the laboratory methodologies used to assess conventional antibacterials are well established, we asseassed their efficacy in evaluating phage plus antibacterial.

**Methods:** 100 multidrug resistant *E. coli* strains were used with three previously isolated and characterised *E. coli* phages of known efficacy. These phages UP17, JK08, 113 were assessed both individually and in a 1:1:1cocktail. In a Phage Microbial Inhibitory Concentration (PmIC) assay, a range of phage concentrations from 10^1^ -10^8^ were inoculated with 5×10^5^/well bacteria in microtitre plates. The first lysed, clear well was taken as the PmIC. Amikacin(AMI) and meropenem(MERO) MICs were determined by microbroth dilution methods(ISO 2776-1:2019) and in combination AMI and MERO MICs were measured with a fixed Phage concentration of 10^5^/well. MICs were performed in triplicate. Time-Kill curves(TKC) were conducted at fosfomycin concentrations of 133, 50 and 5mg/L with and without phage.

**Results:** The PmIC_50/90_ for UP17 were >10^8^/>10^8^; JK08 10^7^/>10^8^; 113 10^7^/>10^8^ and the 1:1:1 cocktail 10^6^/>10^8^. AMI MIC_50/90_ were 0.5/>16 and MERO 0.12/>16mg/L. The addition of UP17 to AMI increased AMI MICs >2 fold in 78 strains. Equivalent increases in AMI MIC were seen with 39 strains with JK08, 54 strains with 113 and 45 strains with the cocktails. In contrast, meropenem MICs in the presence of phage were reduced >2 fold in 24 strains with UP17. Equivalent decreases in MERO MIC were seen with 34 strains with JK08, 26 strains with 113 and 29 strains with the cocktails. In TKCs addition of phage suppressed regrowth.

**Conclusion:** Microbroth methodologies based on ISO 2776-1:2019 and TKCs allow the interaction between Phages and antibacterials to be studied. Optimisation may produce laboratory-based methods with translational value.

## Introduction

Antibiotics have been the primary method of treating bacterial infections since the 1940s due to their ease of administration, safety and potent antibacterial activity (1). This has led to a rapid and ongoing increase in antibiotic resistance despite significant global efforts to reduce the burden of resistance (2). The World Health organisation has stated that antimicrobial resistance (AMR) is one of the top 10 global health threats facing humanity which has forced an interest in alternative anti-infective strategies for the management of bacteria, parasite, viruses, and fungi infections (3).

Bacteriophage were first discovered in 1919 but with the development of antimicrobial drugs within the pharmaceutical industry, the amount of investment in phage standardisation methodologies, commercial phage preparations and regulatory infrastructure was reduced.

Presently, reinvestment and innovation have reignited research in phage studies with the purpose of phage being used for bacteriophage therapy, as an alternative to or in combination with conventional antibiotics (4). Previous studies have shown that single bacteriophage (Phage) strains and Phage cocktails can be used effectively to treat a variety of bacterial infections including those caused by *E.coli* (5). As the mechanism of antibacterial action by phages is completely different from that of antibiotics, this leads to questions as to whether laboratory testing will be needed prior to human use and which methodologies would be optimal in providing reliable and useful therapeutic information. In this regard as there are already laboratory methodologies to test conventional anti-bacterial drugs could phages be tested in the same way.

If phage therapy is to be a viable and widely used in clinical practice, then this should be within (or closely related to) already established antimicrobial testing. We currently have governing bodies which monitor and insure correct antimicrobial usage and susceptibility testing and here we aim to establish if phage and or phage plus antibiotic combinations can be evaluated using these existing methodologies.

## Materials and Methods

### Bacteria, Media and Antibiotics

Bacterial Strains -100 clinical strains were tested from the collection at Southmead hospital. The panel was enriched for resistant strains including, ESBL and carbapenemase (NDM, KPC, OXA-48, IMP and VIM) producers (Supplementary Table S1).

The medium used for MIC and P_M_IC determination was Mueller-Hinton broth II (BD 212322). Media used for phage propagation challenge strains was Mueller Hinton agar (Oxoid PO0152A).

Antibiotics used were Amikacin (Sigma Aldrich) and Meropenem (Biosynth Carbosynth)

### Phage strains, Challenge strains

Phage strains: JK08, 113, UP17 and challenge E coli strains MH10, B31 and EA2 were used. JK08, 113 and UP17 were combined in a cocktail at a ratio of 1:1:1at titres of 1.5 x 10^3^, 10^6^, 10^8^ PFU/mL).

### Phage propagation and titration

Phage propagation and titration - Challenge bacterial strains colonies were inoculated in Mueller Hinton agar (Oxoid PO0152A) and grown overnight at 37^0^C aerobically. To prepare liquid culture, challenge strain colonies were inoculated into LB broth (Oxoid L3147) and grown overnight at 37°C at 100 rpm. Phage propagation was performed by combining phage stains JK08, 113 and UP17 to the appropriate challenge strain at 10^7^ PFU/mL. Bacteria/phage mix cultures were incubated at 37°C at 100 rpm for 6 ± 1 hours. Bacteria/phage mix are centrifuged for 15 mins at 4200×g, and supernatant filtered using (0.2 micron). Phage titre determined by serial dilution (using SM buffer) and plated via plaque assay techniques. Phage titre stored at 4-8°C until use and appropriate dilutions performed as required.

### MIC testing of antimicrobials

Standard MIC determination - ISO 2776-1:2019 and EUCAST Manual V11.00 (Jan 2023) (6,7) were used to determine MICs for two antibiotics. All antibiotic compound and media are stored as per manufacturer instructions. Inoculum preparation was via direct colony suspension method 0.5 McFarland (1.5×10^8^ CFU/mL) and suspension should be used within 60 mins of preparation. Plates were incubated at 36^0^C ±1°C aerobically within 15 mins of inoculation. Plates were read at 18 ± 2 hours. Plates are read manually and EUCAST reading rules are followed (V11.0 – January 2023) and E coli ATCC 25922 QC strains checked to be within acceptable ranges. MIC values interpretation was performed according to current breakpoints (v14.0) Tables at http://www.eucast.org/clinical_breakpoints

### In vitro susceptibility testing Phage Inhibitory Concentration (P_M_IC)

Mueller-Hinton broth II (BD 212322) was used throughout. In all phage titre experiments it was stored prior to use at 4°C and used within 8 hours. The range of phage titres used was 10, 10^2^,10^3^, 10^4^, 10^5^, 10^6^, 10^7^, 10^8^ PFU/mL. Bacterial inoculum preparation was by direct colony suspension (0.5 McFarland approximately 1.5×10^8^ CFU/mL) which was then diluted to give a of 5.5×10^7^ CFU/mL, from which 1µL was added to 100µL of phage suspension, final bacterial inoculum of 5.5×10^5^ CFU/mL. Bacterial suspension was used within 60 mins of preparation. P_M_IC plates were incubated at 36^0^C ±1°C aerobically within 15 mins of inoculation. Plates were manually read at 18 ± 2 hours using EUCAST reading rules. P_M_IC is determined by the first clear well which is directly comparable to standard antimicrobial microbroth MIC testing.

### In vitro susceptibility testing of bacteriophage antimicrobial combinations

A phage titre of 1.0×10^5^ PFU/well was used, added to a doubling dilution series of amikacin or meropenem as for a standard MIC determination. Bacterial inoculum preparation was performed as above, and plates read manually using EUCAST reading rules. ATCC 25922 *<u>E. coli</u>* QC strain used.

### Time Kill curves

Prior to Time Kill curve (TKC) experiments, *E coli* 44913 (fosfomycin MIC 8mg/L) was sub-cultured onto Columbia blood agar (Oxoid PB0123). A 0.5 McFarland inoculum was prepared and diluted to 1.5 x10^6^ CFU/mL in Mueller Hinton II broth (BD 212322) medium. Fosfomycin (Infectopharm) was added a clinically relevant concentrations of 133, 50 and 5mg/L. (AUC24/MIC ratio 400, 150 and 15 respectively) at T0 Post bacterial inoculation. Then a fixed amount of phage cocktail at a titre of 1.5×10^6^ PFU/mL for all fosfomycin concentrations (in a ratio of 1:1:1 of JK08, 113, UP17 respectively) was added at T0 Post bacterial inoculation. Bacterial growth was monitored over a time course of T0 (pre), 1, 2, 3, 4, 5, 6, 24 and 48 hours in triplicate experiments. For every time point 200µL of culture was sampled and plated on Mueller Hinton Agar (Oxoid P01191) to determine CFU enumeration via Don Whitley spiral plater. Inoculated plates are incubated in an aerobic atmosphere for 18-20 hours. Phage PFU was assessed via plaque assay. Area under the bacterial kill curves were calculated as outlined in Noel et al (8).

## Results

### MIC testing of antimicrobials

The MIC_50_, MIC_90_ and range (mg/L) for amikacin was 0.5, >16 and 0.12 to >16mg/L and equivalent values or meropenem was 0.12, >16 and 0.03 to >16mg/L.

### Phage Inhibitory Concentration (P_M_IC) testing

The P_m_IC results for phages UP 17 n=39 strains, JK08 n=41 strains, 113 n=40 and the 1:1:1 cocktail n=41 are shown on Table 1 and individual strain data shown in supplementary materials Table S2.

**Table 1.** Inhibitory Phage titre (PFU/mL) for single phages and the cocktail.

| Bacteriophage | PmlC range | PmlC<sub>50</sub> | PmlC<sub>90</sub> |
| --- | --- | --- | --- |
| UP17 | $10^2 - >10^8$ | $>10^8$ | $>10^8$ |
| JK08 | $10^3 - >10^8$ | $10^7$ | $>10^8$ |
| 113 | $10^2 - >10^8$ | $10^7$ | $>10^8$ |
| 1:1:1 Cocktail | $10^2 - >10^8$ | $10^6$ | $>10^8$ |

The PmIC_50/90_ for UP17 were >10^8^/>10^8^, range 10^2^ to >10^8^ PFU/mL; JK08 10^7^/>10^8^, range 10^3^ to >10^8^; 113 10^7^/>10^8^, range 10^2^ to >10^8^ and the 1:1:1 cocktail 10^6^/>10^8^, range 10^2^ to >10^8^ PFU/mL (Table 1).

Antimicrobial MICs in the presence of phage.

Table 2 shows a comparison of the amikacin MICs alone for 100 *E.coli* strains compared to the amikacin MICs for amikacin in the presence of phages UP17, JK08, 113 and the cocktail. The data are shown graphically in Supplementary Materials Figure S1.

**Table 2.** Comparison of amikacin MICs with and without added bacteriophage.

| Amikacin plus<br>UP17 MIC mg/L | Amikacin<br>MIC mg/L |  |  |  |  |  |  |  |  |
| --- | --- | --- | --- | --- | --- | --- | --- | --- | --- |
|  | ≤0.12 | 0.25 | 0.5 | 1 | 2 | 4 | 8 | ≥16 | Total |
| ≤0.12 |  |  |  |  |  |  |  |  |  |
| 0.25 |  |  |  |  |  |  |  | 1 | 1 |
| 0.5 |  |  | 1 | 1 |  |  |  |  | 2 |
| 1 |  | 2 | 2 |  |  |  |  |  | 4 |
| 2 | 2 | 12 | 4 | 1 | 2 | 1 |  | 2 | 24 |
| 4 | 1 | 12 | 15 | 3 | 1 |  | 1 |  | 33 |
| 8 | 1 | 10 | 2 |  |  |  |  | 2 | 15 |
| ≥16 | 1 | 12 | 4 |  |  |  | 1 | 2 | 21 |
| Total | 5 | 48 | 28 | 5 | 4 | 1 | 2 | 7 | 100 |

| Amikacin plus<br>JK08 MIC mg/L | Amikacin<br>MIC mg/L |  |  |  |  |  |  |  |  |
| --- | --- | --- | --- | --- | --- | --- | --- | --- | --- |
|  | ≤0.12 | 0.25 | 0.5 | 1 | 2 | 4 | 8 | ≥16 | Total |
| ≤0.12 | 2 | 20 | 9 | 2 | 1 | 1 | 1 | 3 | 39 |
| 0.25 |  |  |  |  |  |  |  | 1 | 1 |
| 0.5 |  |  | 1 |  |  |  |  | 1 | 2 |
| 1 | 1 | 3 | 6 | 1 | 1 |  |  |  | 12 |
| 2 | 1 | 10 | 1 | 1 | 2 |  |  |  | 15 |
| 4 | 1 | 8 | 6 | 1 |  |  |  |  | 16 |
| 8 |  | 3 | 2 |  |  |  |  | 2 | 7 |
| ≥16 |  | 4 | 3 |  |  |  | 1 |  | 8 |
| Total | 5 | 48 | 28 | 5 | 4 | 1 | 2 | 7 | 100 |

| Amikacin plus<br>113 MIC mg/L | Amikacin<br>MIC mg/L |  |  |  |  |  |  |  |  |
| --- | --- | --- | --- | --- | --- | --- | --- | --- | --- |
|  | ≤0.12 | 0.25 | 0.5 | 1 | 2 | 4 | 8 | ≥16 | Total |
| ≤0.12 |  | 11 | 6 | 2 | 2 | 1 |  | 2 | 24 |
| 0.25 |  | 1 | 1 |  |  |  |  | 1 | 3 |
| 0.5 |  |  |  |  |  |  | 1 |  | 1 |
| 1 | 1 | 2 | 1 |  |  |  |  |  | 4 |
| 2 | 1 | 10 | 6 | 1 | 1 |  |  | 1 | 20 |
| 4 | 2 | 8 | 7 | 1 | 1 |  |  |  | 19 |
| 8 |  | 5 | 2 |  |  |  |  | 1 | 9 |
| ≥16 | 1 | 11 | 5 |  |  |  | 1 | 2 | 20 |
| Total | 5 | 48 | 28 | 5 | 4 | 1 | 2 | 7 | 100 |

| Amikacin plus<br>1:1:1 cocktail<br>MIC mg/L | Amikacin<br>MIC mg/L |  |  |  |  |  |  |  |  |
| --- | --- | --- | --- | --- | --- | --- | --- | --- | --- |
|  | ≤0.12 | 0.25 | 0.5 | 1 | 2 | 4 | 8 | ≥16 | Total |
| ≤0.12 | 1 | 14 | 10 | 2 | 2 | 1 | 1 | 2 | 33 |
| 0.25 |  |  |  |  |  |  |  | 1 | 1 |
| 0.5 |  | 1 | 2 |  |  |  |  |  | 3 |
| 1 | 1 | 4 | 1 | 1 | 1 |  |  |  | 8 |
| 2 | 2 | 8 | 7 | 1 |  |  |  | 1 | 19 |
| 4 |  | 10 | 2 | 1 | 1 |  |  | 1 | 15 |
| 8 |  | 4 | 4 |  |  |  |  | 1 | 9 |
| ≥16 | 1 | 7 | 2 |  |  |  | 1 | 1 | 12 |
| Total | 5 | 48 | 28 | 5 | 4 | 1 | 2 | 7 | 100 |

Addition of phage UP17 resulted in an increase in amikacin MIC of >2 fold for 78 of 100 strains while addition of JK08 increased MICs >2 fold for 39 strains and 113 increased MICs >2 fold for 54 strains. The cocktail of phages resulted in > 2-fold increases in amikacin MICs for 45 strains. Less than 10 strains showed a >2-fold decrease in amikacin MIC with the addition of UP17, JK08 or the cocktail.

In contrast addition of phage UP17 resulted in a decrease in meropenem MIC of >2 fold for 24 of 100 strains while addition of JK08 decreased MICs >2 fold for 34 strains and 113 decreased MICs >2 fold for 26 strains. The cocktail of phages resulted in >2-fold deceases in meropenem MICs for 29 strains. No strains showed an increase of >2 fold in MIC value (Table 3 and Supplementary Material Figure S2

**Table 3.** Comparison of meropenem MICs with and without added bacteriophage.

| Meropenem<br>plus UP17 MIC<br>mg/L | Meropenem<br>MIC mg/L |  |  |  |  |  |  |  |  |  |
| --- | --- | --- | --- | --- | --- | --- | --- | --- | --- | --- |
|  | ≤0.06 | 0.12 | 0.25 | 0.5 | 1 | 2 | 4 | 8 | ≥16 | Total |
| ≤0.06 | 1 |  |  |  |  |  |  |  |  | 1 |
| 0.12 | 1 | 53 | 7 | 5 | 1 | 1 | 15 | 4 |  | 87 |
| 0.25 |  |  |  |  |  |  |  |  |  | 0 |
| 0.5 |  |  |  |  |  |  |  |  |  | 0 |
| 1 |  |  | 1 |  |  |  |  |  |  | 1 |
| 2 |  |  |  |  |  |  | 1 |  |  | 1 |
| 4 |  |  |  |  |  | 1 | 1 |  |  | 2 |
| 8 |  |  |  |  |  |  |  |  |  | 0 |
| ≥16 |  |  |  |  |  |  |  |  | 8 | 0 |
| Total | 2 | 53 | 8 | 5 | 1 | 2 | 17 | 4 | 8 | 100 |

| Meropenem<br>plus JK08 MIC<br>mg/L | Meropenem<br>MIC mg/L |  |  |  |  |  |  |  |  |  |
| --- | --- | --- | --- | --- | --- | --- | --- | --- | --- | --- |
|  | ≤0.06 | 0.12 | 0.25 | 0.5 | 1 | 2 | 4 | 8 | ≥16 | Total |
| ≤0.06 | 1 | 31 | 6 | 5 | 1 | 1 | 10 | 3 | 2 | 60 |
| 0.12 | 1 | 22 | 2 |  |  |  | 5 | 1 |  | 31 |
| 0.25 |  |  |  |  |  |  |  |  |  | 0 |
| 0.5 |  |  |  |  |  | 1 |  |  |  | 1 |
| 1 |  |  |  |  |  |  |  |  |  | 0 |
| 2 |  |  |  |  |  |  |  |  |  | 0 |
| 4 |  |  |  |  |  |  | 1 |  |  | 1 |
| 8 |  |  |  |  |  |  |  |  |  | 0 |
| ≥16 |  |  |  |  |  |  |  |  | 6 | 6 |
| Total | 2 | 53 | 8 | 5 | 1 | 2 | 16 | 4 | 8 | 99 |

| Meropenem<br>plus 113 MIC<br>mg/L | Meropenem<br>MIC mg/L |  |  |  |  |  |  |  |  |  |
| --- | --- | --- | --- | --- | --- | --- | --- | --- | --- | --- |
|  | ≤0.06 | 0.12 | 0.25 | 0.5 | 1 | 2 | 4 | 8 | ≥16 | Total |
| ≤0.06 |  | 1 |  | 1 |  |  | 1 | 1 |  | 4 |
| 0.12 | 2 | 51 | 7 | 4 | 1 | 1 | 14 | 3 |  | 83 |
| 0.25 |  | 1 | 1 |  |  |  |  |  |  | 2 |
| 0.5 |  |  |  |  |  |  |  |  |  | 0 |
| 1 |  |  |  |  |  |  |  |  |  | 0 |
| 2 |  |  |  |  |  |  | 1 |  |  | 1 |
| 4 |  |  |  |  |  | 1 | 1 |  |  | 2 |
| 8 |  |  |  |  |  |  |  |  |  | 0 |
| ≥16 |  |  |  |  |  |  |  |  | 8 | 8 |
| Total | 2 | 53 | 8 | 5 | 1 | 2 | 17 | 4 | 8 | 100 |

| Meropenem<br>1:1:1 cocktail<br>MIC mg/L | Meropenem<br>MIC mg/L |  |  |  |  |  |  |  |  |  |
| --- | --- | --- | --- | --- | --- | --- | --- | --- | --- | --- |
|  | ≤0.06 | 0.12 | 0.25 | 0.5 | 1 | 2 | 4 | 8 | ≥16 | Total |
| ≤0.06 | 1 | 25 | 6 | 4 |  | 1 | 6 | 2 | 1 | 46 |
| 0.12 | 1 | 27 | 2 |  |  |  | 7 | 2 |  | 39 |
| 0.25 |  | 1 |  |  | 1 |  |  |  |  | 2 |
| 0.5 |  |  |  |  |  | 1 |  |  |  | 1 |
| 1 |  |  |  | 1 |  |  |  |  |  | 1 |
| 2 |  |  |  |  |  |  |  |  |  | 1 |
| 4 |  |  |  |  |  |  | 3 |  | 1 | 4 |
| 8 |  |  |  |  |  |  | 1 |  |  | 1 |
| ≥16 |  |  |  |  |  |  |  |  | 6 | 6 |
| Total | 2 | 53 | 8 | 5 | 1 | 2 | 17 | 4 | 8 | 100 |

### Time-Kill Curves

Figure 1 shows the time-kill curves for fosfomycin at three concentrations with the presence of phage compared to fosfomycin alone and phage alone. Phage alone produced a 1-2log reduction in bacterial count by 6hrs while the addition of phage to fosfomycin reduced viable bacterial counts at either 24h or 48h incubation. Areas-under-the -bacterial-kill curves (AUBKC) were calculated and are shown in Supplementary Material Table S3 – fosfomycin plus phage produced smaller AUBKC values than fosfomycin alone at 24h and 48h showing the additional suppression of bacterial burden by the addition of phage.

**Figure 1.**
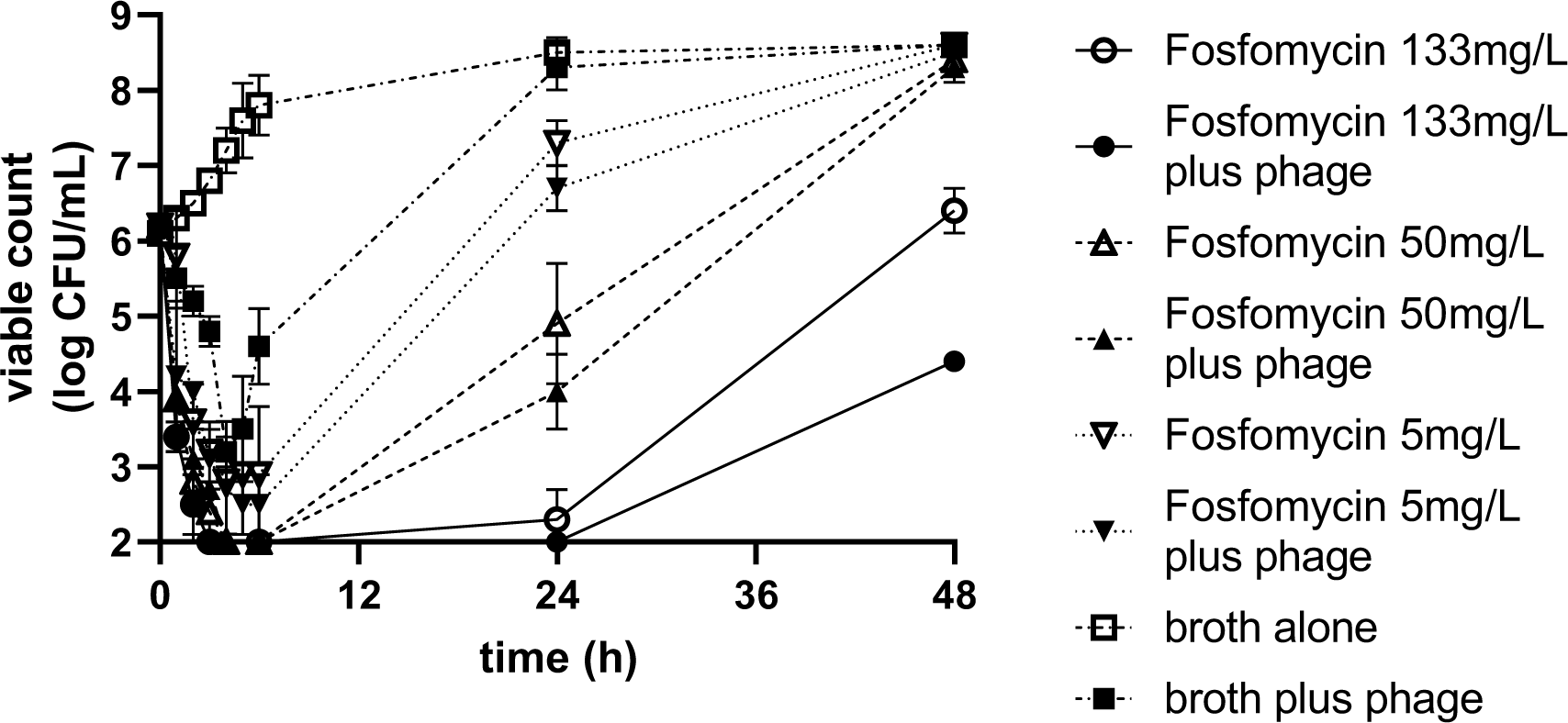
Time kill curves for Fosfomycin at high, medium and low concentratios in the presence of phage cocktail.

## Conclusions

The *in vitro* protocols used here, which are based on those used to assess existing antibacterials, have been shown to be an effective and reproducible methods of determining P_M_IC, antibacterial MICs in the presence of bacteriophage and also in time-kill curves. However, it is also important to note the validation work in this study was primarily for the laboratory techniques used rather than to evaluate the probable clinical value of using single or multiple phage strains in combination with antimicrobials. Most anti-bacterial and phage combination assessment is based on antibacterial synergy combination methods resulting in FIC calculations (8,9) which may be helpful in highly specific strain interactions, but this does have limitations in terms of screening purposes and also provides poorly translatable information. Pharmacodynamic thinking related to antibacterial combinations would indicate that total antibacterial activity is more important than arbitrary definitions of synergy or antagonism in providing useful translational information (10).

The combination of amikacin with phages produced a different pattern of interaction than that of meropenem plus phage in the PmIC determinations. Amikacin combinations tended to result in an increase in amikacin MICs by >2 fold with a minority of strains showing a decrease in MIC of >2 fold. In contrast the combination of meropenem plus phages was more likely to produce a reduction of MIC of >2 fold and in no strains was the MICs increased >2 fold. Differences in the interaction between phages and different classes of antibiotics has been described before using more complex and less translatable technology (9). In addition, it has been shown that phage plus B lactams produce different morphological changes in *E coli* when compared to phage plus amikacin.

Time -kill curves have been used previously to study the impact of combining phages and antimicrobials but again using concepts of synergy and antagonism and like here using static antimicrobial concentrations (12). The ability to study phage plus antimicrobial combinations in static concentration systems is an initial step to using more complex in vitro tools which can simulate dynamically changing concentrations of phage and/or antibiotic which in turn may enable more clinically valuable information to be assembled (13).

This is encouraging data; it indicates existing methodologies can be used to assess phages and phages plus antibacterial combinations. However, there is much more development work to do to show how such methodologies can be applied to clinical problems and understand any potential limitations. This should include studying the effect of variation in the phage titre used, alterations of phage cocktail component ratio, the use of other antimicrobial classes plus phage cocktail combinations, evaluating the timing of combining phages and antimicrobials and any impacts on bacterial resistance to phages and/or antibiotics. In addition, the *in vivo* predictive value of such *in vitro* tests should be assessed in animal infection models and the more detailed pharmacodynamics of phages scrutinised both *in vitro* and *in vivo* as many questions remain unresolved (14)

## Transparency Declarations

APM holds research grants/activities with Merck, Shionogi, InfectoPharm, GSK, Roche, Bioversys, and Nosopharm. MA and ARN declare no competing interests.

## Supplementary material

**Table S1).** E coli strains used in bacteriophage and antibiotic testing.

| Number | Panel number | Category | Maldi ID | Maldi Score |
| --- | --- | --- | --- | --- |
| 1 | C1.1 | Quinolone S | *Escherichia coli* | 2.31 |
| 2 | C1.2 | Quinolone S | *Escherichia coli* | 2.43 |
| 3 | C1.3 | Quinolone S | *Escherichia coli* | 2.46 |
| 4 | C1.4 | Quinolone S | *Escherichia coli* | 2.36 |
| 5 | C1.5 | Quinolone S | *Escherichia coli* | 2.29 |
| 6 | C1.6 | Quinolone S | *Escherichia coli* | 2.05 |
| 7 | C1.7 | Quinolone S | *Escherichia coli* | 2.24 |
| 8 | C1.8 | Quinolone S | *Escherichia coli* | 2.24 |
| 9 | C1.9 | Quinolone S | *Escherichia coli* | 2.26 |
| 10 | C1.10 | Quinolone S | *Escherichia coli* | 2.39 |
| 11 | C1.11 | Quinolone S | *Escherichia coli* | 2.5 |
| 12 | C1.12 | Quinolone S | *Escherichia coli* | 2.21 |
| 13 | C1.13 | Quinolone S | *Escherichia coli* | 2 |
| 14 | C1.14 | Quinolone S | *Escherichia coli* | 2.38 |
| 15 | C1.15 | Quinolone S | *Escherichia coli* | 2.33 |
| 16 | C1.16 | Quinolone S | *Escherichia coli* | 2.3 |
| 17 | C1.17 | Quinolone S | *Escherichia coli* | 2.47 |
| 18 | C1.18 | Quinolone S | *Escherichia coli* | 2.17 |
| 19 | C1.19 | Quinolone S | *Escherichia coli* | 2.25 |
| 20 | C1.20 | Quinolone S | *Escherichia coli* | 2.39 |
| 21 | C1.21 | R Quinolone | *Escherichia coli* | 2.18 |
| 22 | C1.22 | R Quinolone | *Escherichia coli* | 2.27 |
| 23 | C1.23 | R Quinolone | *Escherichia coli* | 2.36 |
| 24 | C1.24 | R Quinolone | *Escherichia coli* | 2.25 |
| 25 | C1.25 | R Quinolone | *Escherichia coli* | 2.23 |
| 26 | C1.26 | R Quinolone | *Escherichia coli* | 2.32 |
| 27 | C1.27 | R Quinolone | *Escherichia coli* | 2.29 |
| 28 | C1.28 | R Quinolone | *Escherichia coli* | 2.22 |
| 29 | C1.29 | R Quinolone | *Escherichia coli* | 2.25 |
| 30 | C1.30 | R Quinolone | *Escherichia coli* | 2.17 |
| 31 | C1.31 | R Quinolone | *Escherichia coli* | 2.19 |
| 32 | C1.32 | R Quinolone | *Escherichia coli* | 2.27 |
| 33 | C1.33 | R Quinolone | *Escherichia coli* | 2.28 |
| 34 | C1.34 | R Quinolone | *Escherichia coli* | 2.03 |
| 35 | C1.35 | R Quinolone | *Escherichia coli* | 2.37 |
| 36 | C1.36 | R Quinolone | *Escherichia coli* | 2.43 |
| 37 | C1.37 | R Quinolone | *Escherichia coli* | 2.25 |
| 38 | C1.38 | R Quinolone | *Escherichia coli* | 2.33 |
| 39 | C1.39 | R Quinolone | *Escherichia coli* | 2.17 |
| 40 | C1.40 | R Quinolone | *Escherichia coli* | 2.14 |
| 41 | C1.41 | ESBL | *Escherichia coli* | 2.29 |
| 42 | C1.42 | ESBL | *Escherichia coli* | 2.33 |
| 43 | C1.43 | ESBL | *Escherichia coli* | 2.4 |
| 44 | C1.44 | ESBL | *Escherichia coli* | 2.41 |
| 45 | C1.45 | ESBL | *Escherichia coli* | 2.32 |
| 46 | C1.46 | ESBL | *Escherichia coli* | 2.36 |
| 47 | C1.47 | ESBL | *Escherichia coli* | 2.14 |
| 48 | C1.48 | ESBL | *Escherichia coli* | 2.17 |
| 49 | C1.49 | ESBL | *Escherichia coli* | 2.29 |
| 50 | C1.50 | ESBL | *Escherichia coli* | 2.33 |
| 51 | C1.51 | ESBL | *Escherichia coli* | 2.37 |
| 52 | C1.52 | ESBL | *Escherichia coli* | 2.26 |
| 53 | C1.53 | ESBL | *Escherichia coli* | 2.25 |
| 54 | C1.54 | ESBL | *Escherichia coli* | 2.07 |
| 55 | C1.55 | ESBL | *Escherichia coli* | 2.32 |
| 56 | C1.56 | ESBL | *Escherichia coli* | 2.36 |
| 57 | C1.57 | ESBL | *Escherichia coli* | 2.29 |
| 58 | C1.58 | ESBL | *Escherichia coli* | 2.31 |
| 59 | C1.59 | ESBL | *Escherichia coli* | 2.4 |
| 60 | C1.60 | CARBR | *Escherichia coli* | 2.06 |
| 61 | C1.61 | CARBR | *Escherichia coli* | 2.16 |
| 62 | C1.62 | CARBR | *Escherichia coli* | 2.39 |
| 63 | C1.63 | CARBR | *Escherichia coli* | 2.29 |
| 64 | C1.64 | CARBR | *Escherichia coli* | 2.22 |
| 65 | C1.65 | CARBR | *Escherichia coli* | 2.52 |
| 66 | C1.66 | CARBR | *Escherichia coli* | 2.35 |
| 67 | C1.67 | CARBR | *Escherichia coli* | 2.23 |
| 68 | C1.68 | CARBR | *Escherichia coli* | 2.37 |
| 69 | C1.69 | CARBR | *Escherichia coli* | 2.33 |
| 70 | C1.70 | N/S MDR | *Escherichia coli* | 2.29 |
| 71 | C1.71 | N/S MDR | *Escherichia coli* | 2.2 |
| 72 | C1.72 | N/S MDR | *Escherichia coli* | 2.33 |
| 73 | C1.73 | N/S MDR | *Escherichia coli* | 2.26 |
| 74 | C1.74 | N/S MDR | *Escherichia coli* | 2.2 |
| 75 | C1.75 | N/S MDR | *Escherichia coli* | 2.24 |
| 76 | C1.76 | N/S MDR | *Escherichia coli* | 2.28 |
| 77 | C1.77 | N/S MDR | *Escherichia coli* | 2.33 |
| 78 | C1.78 | N/S MDR | *Escherichia coli* | 2.29 |
| 79 | C1.79 | N/S MDR | *Escherichia coli* | 2.03 |
| 80 | C1.80 | N/S MDR | *Escherichia coli* | 2.2 |
| 81 | C1.81 | N/S MDR | *Escherichia coli* | 2.31 |
| 82 | C1.82 | N/S MDR | *Escherichia coli* | 2.14 |
| 83 | C1.83 | N/S MDR | *Escherichia coli* | 2.38 |
| 84 | C1.84 | N/S MDR | *Escherichia coli* | 2.38 |
| 85 | C1.85 | N/S MDR | *Escherichia coli* | 2.34 |
| 86 | C1.86 | N/S MDR | *Escherichia coli* | 2.37 |
| 87 | C1.87 | N/S MDR | *Escherichia coli* | 2.28 |
| 88 | C1.88 | N/S MDR | *Escherichia coli* | 2.27 |
| 89 | C1.89 | N/S MDR | *Escherichia coli* | 2.25 |
| 90 | C1.90 | N/S MDR | *Escherichia coli* | 2.4 |
| 91 | C1.91 | N/S MDR | *Escherichia coli* | 2.39 |
| 92 | C1.92 | N/S MDR | *Escherichia coli* | 2.18 |
| 93 | C1.93 | N/S MDR | *Escherichia coli* | 2.24 |
| 94 | C1.94 | N/S MDR | *Escherichia coli* | 2.26 |
| 95 | C1.95 | N/S MDR | *Escherichia coli* | 2.4 |
| 96 | C1.96 | N/S MDR | *Escherichia coli* | 2.36 |
| 97 | C1.97 | N/S MDR | *Escherichia coli* | 2.13 |
| 98 | C1.98 | N/S MDR | *Escherichia coli* | 2.11 |
| 99 | C1.99 | N/S MDR | *Escherichia coli* | 2.32 |
| 100 | C1.100 | N/S MDR | *Escherichia coli* | 2.03 |

**Table S2).** Line listing P_m_IC results for each E coli strain tested with UP17, 113, JK 08 and 1:1:1 cocktail in triplicate.

|  | UP17 |  |  |  |  |  | UP17 |  |  |  |  |
| --- | --- | --- | --- | --- | --- | --- | --- | --- | --- | --- | --- |
|  | <u>1</u> | <u>2</u> | <u>3</u> | <u>Average</u> | <u>SD</u> |  | <u>1</u> | <u>2</u> | <u>3</u> | <u>Average</u> | <u>SD</u> |
| C1.1 | 1.00E+07 | 1.00E+07 | 1.00E+07 | 1.00E+07 | 0 | C1.41 | 1.00E+08 | 1.00E+08 | 1.00E+08 | 1.00E+08 | 0 |
| C1.2 | 1.00E+04 | 1.00E+04 | 1.00E+04 | 1.00E+04 | 0 | C1.42 | >1.E+08 | >1.E+08 | >1.E+08 | >1.E+08 | 0 |
| C1.3 | 1.00E+08 | 1.00E+08 | 1.00E+08 | 1.00E+08 | 0 | C1.43 | >1.E+08 | >1.E+08 | >1.E+08 | >1.E+08 | 0 |
| C1.4 | 1.00E+06 | 1.00E+06 | 1.00E+06 | 1.00E+06 | 0 | C1.44 | >1.E+08 | >1.E+08 | >1.E+08 | >1.E+08 | 0 |
| C1.5 | 1.00E+06 | 1.00E+06 | 1.00E+06 | 1.00E+06 | 0 | C1.45 | 1.00E+05 | 1.00E+05 | 1.00E+05 | 1.00E+05 | 0 |
| C1.6 | >1.E+08 | >1.E+08 | >1.E+08 | >1.E+08 | 0 | C1.46 | 1.00E+08 | 1.00E+08 | 1.00E+08 | 1.00E+08 | 0 |
| C1.7 | 1.00E+07 | 1.00E+07 | 1.00E+07 | 1.00E+07 | 0 | C1.47 | 1.00E+08 | 1.00E+08 | 1.00E+08 | 1.00E+08 | 0 |
| C1.8 | 1.00E+07 | 1.00E+07 | 1.00E+07 | 1.00E+08 | 0 | C1.48 | 1.00E+07 | 1.00E+07 | 1.00E+07 | 1.00E+07 | 0 |
| C1.9 | 1.00E+04 | 1.00E+04 | 1.00E+04 | 1.00E+04 | 0 | C1.49 | 1.00E+08 | 1.00E+08 | 1.00E+08 | 1.00E+08 | 0 |
| C1.10 | 1.00E+08 | 1.00E+08 | 1.00E+08 | 1.00E+08 | 0 | C1.50 | >1.E+08 | >1.E+08 | >1.E+08 | >1.E+08 | 0 |
| C1.11 | 1.00E+07 | 1.00E+07 | 1.00E+07 | 1.00E+07 | 0 | C1.51 | 1.00E+08 | 1.00E+08 | 1.00E+08 | 1.00E+08 | 0 |
| C1.12 | 1.00E+04 | 1.00E+04 | 1.00E+04 | 1.00E+04 | 0 | C1.52 | 1.00E+08 | 1.00E+08 | 1.00E+08 | 1.00E+08 | 0 |
| C1.13 | >1.E+08 | >1.E+08 | >1.E+08 | >1.E+08 | 0 | C1.53 | 1.00E+08 | 1.00E+08 | 1.00E+08 | 1.00E+08 | 0 |
| C1.14 | >1.E+08 | >1.E+08 | >1.E+08 | >1.E+08 | 0 | C1.54 | 1.00E+08 | 1.00E+08 | 1.00E+08 | 1.00E+08 | 0 |
| C1.15 | >1.E+08 | >1.E+08 | >1.E+08 | >1.E+08 | 0 | C1.55 | >1.E+08 | >1.E+08 | >1.E+08 | >1.E+08 | 0 |
| C1.16 | 1.00E+08 | 1.00E+08 | 1.00E+08 | 1.00E+08 | 0 | C1.56 | >1.E+08 | >1.E+08 | >1.E+08 | >1.E+08 | 0 |
| C1.17 | >1.E+08 | >1.E+08 | >1.E+08 | >1.E+08 | 0 | C1.57 | >1.E+08 | >1.E+08 | >1.E+08 | >1.E+08 | 0 |
| C1.18 | >1.E+08 | >1.E+08 | >1.E+08 | >1.E+08 | 0 | C1.58 | >1.E+08 | >1.E+08 | >1.E+08 | >1.E+08 | 0 |
| C1.19 | 1.00E+08 | 1.00E+08 | 1.00E+08 | 1.00E+08 | 0 | C1.59 | >1.E+08 | >1.E+08 | >1.E+08 | >1.E+08 | 0 |
| C1.20 | >1.E+08 | >1.E+08 | >1.E+08 | >1.E+08 | 0 |  |  |  |  |  |  |

|  | 113 |  |  |  |  |  | 113 |  |  |  |  |
| --- | --- | --- | --- | --- | --- | --- | --- | --- | --- | --- | --- |
|  | <u>1</u> | <u>2</u> | <u>3</u> | <u>Average</u> | <u>SD</u> |  | <u>1</u> | <u>2</u> | <u>3</u> | <u>Average</u> | <u>SD</u> |
| C1.1 | 1.00E+08 | 1.00E+08 | 1.00E+08 | 1.00E+08 | 0 | C1.81 | 1.00E+03 | 1.00E+03 | 1.00E+03 | 1.00E+03 | 0 |
| C1.2 | 1.00E+04 | 1.00E+04 | 1.00E+04 | 1.00E+04 | 0 | C1.82 | 1.00E+07 | 1.00E+07 | 1.00E+07 | 1.00E+07 | 0 |
| C1.3 | 1.00E+06 | 1.00E+06 | 1.00E+06 | 1.00E+06 | 0 | C1.83 | 1.00E+08 | 1.00E+08 | 1.00E+08 | 1.00E+08 | 0 |
| C1.4 | 1.00E+03 | 1.00E+03 | 1.00E+03 | 1.00E+03 | 0 | C1.84 | >1.E+08 | >1.E+08 | >1.E+08 | >1.E+08 | 0 |
| C1.5 | 1.00E+08 | 1.00E+08 | 1.00E+08 | 1.00E+08 | 0 | C1.85 | >1.E+08 | >1.E+08 | >1.E+08 | >1.E+08 | 0 |
| C1.6 | >1.E+08 | >1.E+08 | >1.E+08 | >1.E+08 | 0 | C1.86 | 1.00E+08 | 1.00E+08 | 1.00E+08 | 1.00E+08 | 0 |
| C1.7 | 1.00E+07 | 1.00E+07 | 1.00E+07 | 1.00E+07 | 0 | C1.87 | >1.E+08 | >1.E+08 | >1.E+08 | >1.E+08 | 0 |
| C1.8 | 1.00E+08 | 1.00E+08 | 1.00E+08 | 1.00E+08 | 0 | C1.88 | 1.00E+08 | 1.00E+08 | 1.00E+08 | 1.00E+08 | 0 |
| C1.9 | 1.00E+02 | 1.00E+02 | 1.00E+02 | 1.00E+02 | 0 | C1.89 | 1.00E+08 | 1.00E+08 | 1.00E+08 | 1.00E+08 | 0 |
| C1.10 | 1.00E+05 | 1.00E+05 | 1.00E+05 | 1.00E+05 | 0 | C1.90 | 1.00E+07 | 1.00E+07 | 1.00E+07 | 1.00E+07 | 0 |
| C1.11 | >1.E+08 | >1.E+08 | >1.E+08 | >1.E+08 | 0 | C1.91 | >1.E+08 | >1.E+08 | >1.E+08 | >1.E+08 | 0 |
| C1.12 | 1.00E+03 | 1.00E+03 | 1.00E+03 | 1.00E+03 | 0 | C1.92 | 1.00E+06 | 1.00E+06 | 1.00E+06 | 1.00E+06 | 0 |
| C1.13 | >1.E+08 | >1.E+08 | >1.E+08 | >1.E+08 | 0 | C1.93 | 1.00E+07 | 1.00E+07 | 1.00E+07 | 1.00E+07 | 0 |
| C1.14 | >1.E+08 | >1.E+08 | >1.E+08 | >1.E+08 | 0 | C1.94 | 1.00E+06 | 1.00E+06 | 1.00E+06 | 1.00E+06 | 0 |
| C1.15 | 1.00E+07 | 1.00E+07 | 1.00E+07 | 1.00E+07 | 0 | C1.95 | 1.00E+08 | 1.00E+08 | 1.00E+08 | 1.00E+08 | 0 |
| C1.16 | 1.00E+08 | 1.00E+08 | 1.00E+08 | 1.00E+08 | 0 | C1.96 | >1.E+08 | >1.E+08 | >1.E+08 | >1.E+08 | 0 |
| C1.17 | 1.00E+06 | 1.00E+06 | 1.00E+06 | 1.00E+06 | 0 | C1.97 | 1.00E+07 | 1.00E+07 | 1.00E+07 | 1.00E+07 | 0 |
| C1.18 | 1.00E+04 | 1.00E+04 | 1.00E+04 | 1.00E+04 | 0 | C1.98 | 1.00E+07 | 1.00E+07 | 1.00E+07 | 1.00E+07 | 0 |
| C1.19 | 1.00E+06 | 1.00E+06 | 1.00E+06 | 1.00E+06 | 0 | C1.99 | >1.E+08 | >1.E+08 | >1.E+08 | >1.E+08 | 0 |
| C1.20 | >1.E+08 | >1.E+08 | >1.E+08 | >1.E+08 | 0 | C1.100 | 1.00E+07 | 1.00E+07 | 1.00E+07 | 1.00E+07 | 0 |

|  | JK08 |  |  |  |  |  | JK08 |  |  |  |  |
| --- | --- | --- | --- | --- | --- | --- | --- | --- | --- | --- | --- |
|  | <u>1</u> | <u>2</u> | <u>3</u> | <u>Average</u> | <u>SD</u> |  | <u>1</u> | <u>2</u> | <u>3</u> | <u>Average</u> | <u>SD</u> |
| C1.21 | 1.00E+05 | 1.00E+05 | 1.00E+05 | 1.00E+05 | 0 | C1.60 | >1.E+08 | >1.E+08 | >1.E+08 | >1.E+08 | 0 |
| C1.22 | 1.00E+05 | 1.00E+05 | 1.00E+05 | 1.00E+05 | 0 | C1.61 | 1.00E+06 | 1.00E+06 | 1.00E+06 | 1.00E+06 | 0 |
| C1.23 | <1.E+01 | <1.E+01 | <1.E+01 | <1.E+01 | 0 | C1.62 | 1.00E+07 | 1.00E+07 | 1.00E+07 | 1.00E+07 | 0 |
| C1.24 | 1.00E+03 | 1.00E+03 | 1.00E+03 | 1.00E+03 | 0 | C1.63 | >1.E+08 | >1.E+08 | >1.E+08 | >1.E+08 | 0 |
| C1.25 | 1.00E+06 | 1.00E+06 | 1.00E+06 | 1.00E+06 | 0 | C1.64 | >1.E+08 | >1.E+08 | >1.E+08 | >1.E+08 | 0 |
| C1.26 | >1.E+08 | >1.E+08 | >1.E+08 | >1.E+08 | 0 | C1.65 | >1.E+08 | >1.E+08 | >1.E+08 | >1.E+08 | 0 |
| C1.27 | 1.00E+07 | 1.00E+07 | 1.00E+07 | 1.00E+07 | 0 | C1.66 | >1.E+08 | >1.E+08 | >1.E+08 | >1.E+08 | 0 |
| C1.28 | 1.00E+07 | 1.00E+07 | 1.00E+07 | 1.00E+07 | 0 | C1.67 | 1.00E+03 | 1.00E+03 | 1.00E+03 | 1.00E+03 | 0 |
| C1.29 | 1.00E+05 | 1.00E+05 | 1.00E+05 | 1.00E+05 | 0 | C1.68 | 1.00E+08 | 1.00E+08 | 1.00E+08 | 1.00E+08 | 0 |
| C1.30 | 1.00E+08 | 1.00E+08 | 1.00E+08 | 1.00E+08 | 0 | C1.69 | 1.00E+08 | 1.00E+08 | 1.00E+08 | 1.00E+08 | 0 |
| C1.31 | >1.E+08 | >1.E+08 | >1.E+08 | >1.E+08 | 0 | C1.70 | 1.00E+06 | 1.00E+06 | 1.00E+06 | 1.00E+06 | 0 |
| C1.32 | 1.00E+05 | 1.00E+05 | 1.00E+05 | 1.00E+05 | 0 | C1.71 | 1.00E+04 | 1.00E+04 | 1.00E+04 | 1.00E+04 | 0 |
| C1.33 | 1.00E+08 | 1.00E+08 | 1.00E+08 | 1.00E+08 | 0 | C1.72 | 1.00E+03 | 1.00E+03 | 1.00E+03 | 1.00E+03 | 0 |
| C1.34 | 1.00E+05 | 1.00E+05 | 1.00E+05 | 1.00E+05 | 0 | C1.73 | 1.00E+08 | 1.00E+08 | 1.00E+08 | 1.00E+08 | 0 |
| C1.35 | 1.00E+06 | 1.00E+06 | 1.00E+06 | 1.00E+06 | 0 | C1.74 | 1.00E+08 | 1.00E+08 | 1.00E+08 | 1.00E+08 | 0 |
| C1.36 | >1.E+08 | >1.E+08 | >1.E+08 | >1.E+08 | 0 | C1.75 | >1.E+08 | >1.E+08 | >1.E+08 | >1.E+08 | 0 |
| C1.37 | 1.00E+03 | 1.00E+03 | 1.00E+03 | 1.00E+03 | 0 | C1.76 | >1.E+08 | >1.E+08 | >1.E+08 | >1.E+08 | 0 |
| C1.38 | 1.00E+07 | 1.00E+07 | 1.00E+07 | 1.00E+07 | 0 | C1.77 | 1.00E+05 | 1.00E+05 | 1.00E+05 | 1.00E+05 | 0 |
| C1.39 | >1.E+08 | >1.E+08 | >1.E+08 | >1.E+08 | 0 | C1.78 | 1.00E+06 | 1.00E+06 | 1.00E+06 | 1.00E+06 | 0 |
| C1.40 | 1.00E+05 | 1.00E+05 | 1.00E+05 | 1.00E+05 | 0 | C1.79 | 1.00E+06 | 1.00E+06 | 1.00E+06 | 1.00E+06 | 0 |
|  |  |  |  |  |  | C1.80 | >1.E+08 | >1.E+08 | >1.E+08 | >1.E+08 | 0 |

|  | 01:01:01 |  |  |  |  |  | 01:01:01 |  |  |  |  |
| --- | --- | --- | --- | --- | --- | --- | --- | --- | --- | --- | --- |
|  | 1 | 2 | 3 | Average | SD |  | 1 | 2 | 3 | Average | SD |
| C1.2 | 1.00E+05 | 1.00E+05 | 1.00E+05 | 1.00E+05 | 0 | C1.61 | 1.00E+05 | 1.00E+05 | 1.00E+05 | 1.00E+05 | 0 |
| C1.5 | 1.00E+04 | 1.00E+04 | 1.00E+04 | 1.00E+04 | 0 | C1.62 | 1.00E+06 | 1.00E+06 | 1.00E+06 | 1.00E+06 | 0 |
| C1.9 | 1.00E+02 | 1.00E+02 | 1.00E+02 | 1.00E+02 | 0 | C1.65 | >1.E+08 | >1.E+08 | >1.E+08 | >1.E+08 | 0 |
| C1.11 | 1.00E+07 | 1.00E+07 | 1.00E+07 | 1.00E+07 | 0 | C1.68 | 1.00E+05 | 1.00E+05 | 1.00E+05 | 1.00E+05 | 0 |
| C1.14 | >1.E+08 | >1.E+08 | >1.E+08 | >1.E+08 | 0 | C1.71 | 1.00E+04 | 1.00E+04 | 1.00E+04 | 1.00E+04 | 0 |
| C1.17 | 1.00E+05 | 1.00E+05 | 1.00E+05 | 1.00E+05 | 0 | C1.74 | 1.00E+08 | 1.00E+08 | 1.00E+08 | 1.00E+08 | 0 |
| C1.19 | 1.00E+05 | 1.00E+05 | 1.00E+05 | 1.00E+05 | 0 | C1.78 | 1.00E+06 | 1.00E+06 | 1.00E+06 | 1.00E+06 | 0 |
| C1.24 | 1.00E+04 | 1.00E+04 | 1.00E+04 | 1.00E+04 | 0 | C1.83 | 1.00E+07 | 1.00E+07 | 1.00E+07 | 1.00E+07 | 0 |
| C1.28 | 1.00E+04 | 1.00E+04 | 1.00E+04 | 1.00E+04 | 0 | C1.86 | 1.00E+06 | 1.00E+06 | 1.00E+06 | 1.00E+06 | 0 |
| C1.32 | 1.00E+06 | 1.00E+06 | 1.00E+06 | 1.00E+06 | 0 | C1.89 | 1.00E+07 | 1.00E+07 | 1.00E+07 | 1.00E+07 | 0 |
| C1.34 | 1.00E+06 | 1.00E+06 | 1.00E+06 | 1.00E+06 | 0 | C1.90 | 1.00E+02 | 1.00E+02 | 1.00E+02 | 1.00E+02 | 0 |
| C1.36 | >1.E+08 | >1.E+08 | >1.E+08 | >1.E+08 | 0 | C1.92 | 1.00E+05 | 1.00E+05 | 1.00E+05 | 1.00E+05 | 0 |
| C1.39 | 1.00E+08 | 1.00E+08 | 1.00E+08 | 1.00E+08 | 0 | C1.93 | 1.00E+06 | 1.00E+06 | 1.00E+06 | 1.00E+06 | 0 |
| C1.42 | >1.E+08 | >1.E+08 | >1.E+08 | >1.E+08 | 0 | C1.94 | 1.00E+06 | 1.00E+06 | 1.00E+06 | 1.00E+06 | 0 |
| C1.45 | 1.00E+02 | 1.00E+02 | 1.00E+02 | 1.00E+02 | 0 | C1.95 | 1.00E+06 | 1.00E+06 | 1.00E+06 | 1.00E+06 | 0 |
| C1.47 | 1.00E+05 | 1.00E+05 | 1.00E+05 | 1.00E+05 | 0 | C1.96 | >1.E+08 | >1.E+08 | >1.E+08 | >1.E+08 | 0 |
| C1.48 | 1.00E+05 | 1.00E+05 | 1.00E+05 | 1.00E+05 | 0 | C1.97 | 1.00E+06 | 1.00E+06 | 1.00E+06 | 1.00E+06 | 0 |
| C1.51 | 1.00E+08 | 1.00E+08 | 1.00E+08 | 1.00E+08 | 0 | C1.98 | 1.00E+07 | 1.00E+07 | 1.00E+07 | 1.00E+07 | 0 |
| C1.53 | 1.00E+08 | 1.00E+08 | 1.00E+08 | 1.00E+08 | 0 | C1.99 | >1.E+08 | >1.E+08 | >1.E+08 | >1.E+08 | 0 |
| C1.54 | 1.00E+08 | 1.00E+08 | 1.00E+08 | 1.00E+08 | 0 | C1.100 | 1.00E+06 | 1.00E+06 | 1.00E+06 | 1.00E+06 | 0 |
| C1.57 | 1.00E+08 | 1.00E+08 | 1.00E+08 | 1.00E+08 | 0 |  |  |  |  |  |  |

**Table S3).** Area-under-the-bacterial-Kill-curves (AUBKC) for fosfomycin alone, phage cocktail alone and combinations of phage and fosfomycin.

|  | AUBKC (logCFU/mL.h) |  |  |
| --- | --- | --- | --- |
|  | 0-6hr | 0-24hr | 0-48hr |
| Fosfomycin 133mg/L | 4.0±0.6 | 6.5±4.0 | 62.7±5.7 |
| Fosfomycin 133mg/L + phage | 4.0± 0.7 | 6.4±3.1 | 38.2±9.2 |
| Fosfomycin 50mg/L | 5.2±1.0 | 31.0±6.4 | 142.8±18.8 |
| Fosfomycin 50mg/L + phage | 5.8± 0.5 | 23.9±3.7 | 124.2±11.7 |
| Fosfomycin 5mg/L | 10.8±0.7 | 66.6±8.7 | 209.9±10.8 |
| Fosfomycin 5mg/L + phage | 8.8±2.3 | 56.1±6.5 | 191.2±8.9 |
| Phage alone | 15.6±1.7 | 96.2±6.6 | 251.1±9.4 |
| Growth Control | 29.8±0.8 | 139.1±3.7 | 295.8±2.9 |

**Figure S1).**
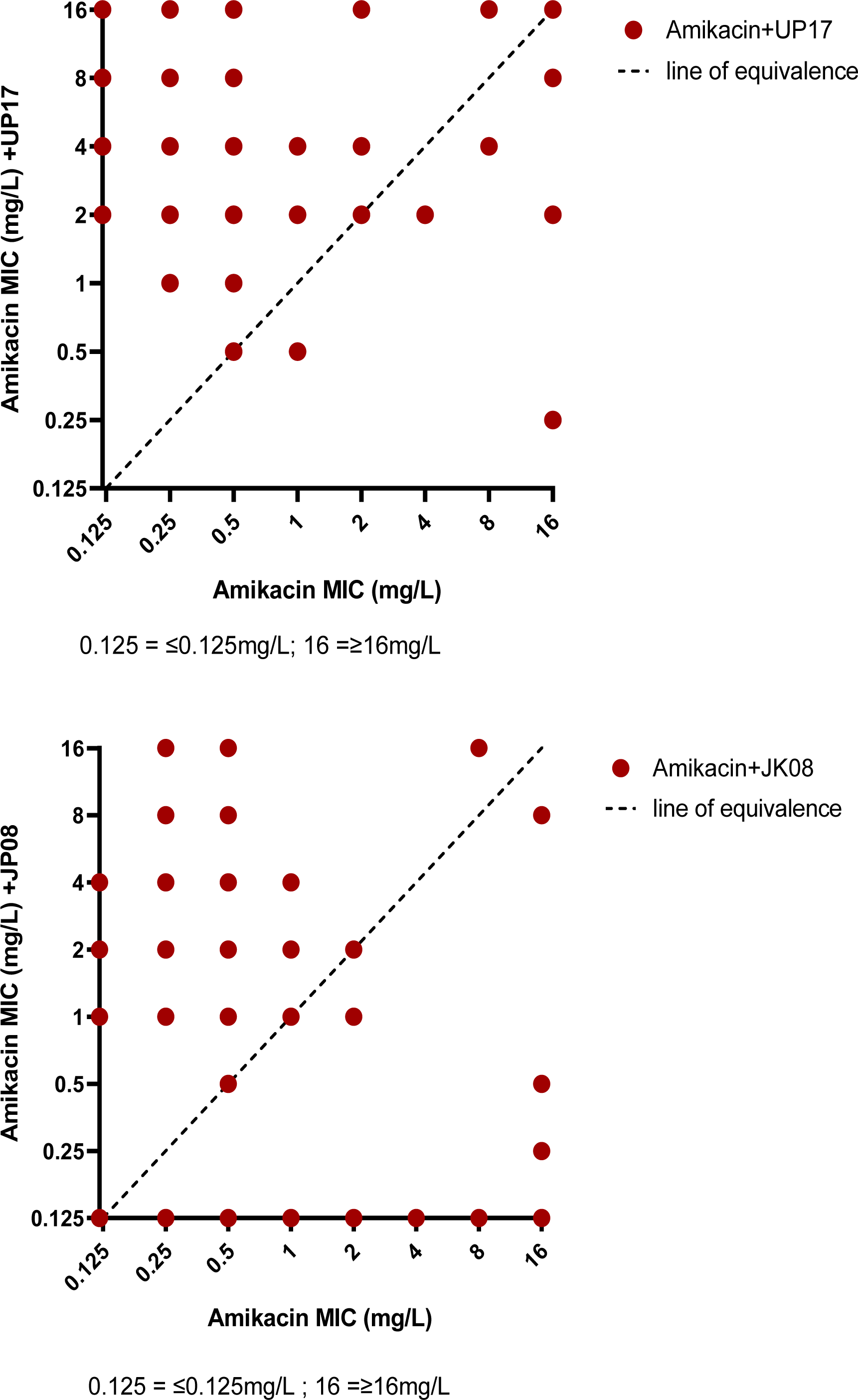
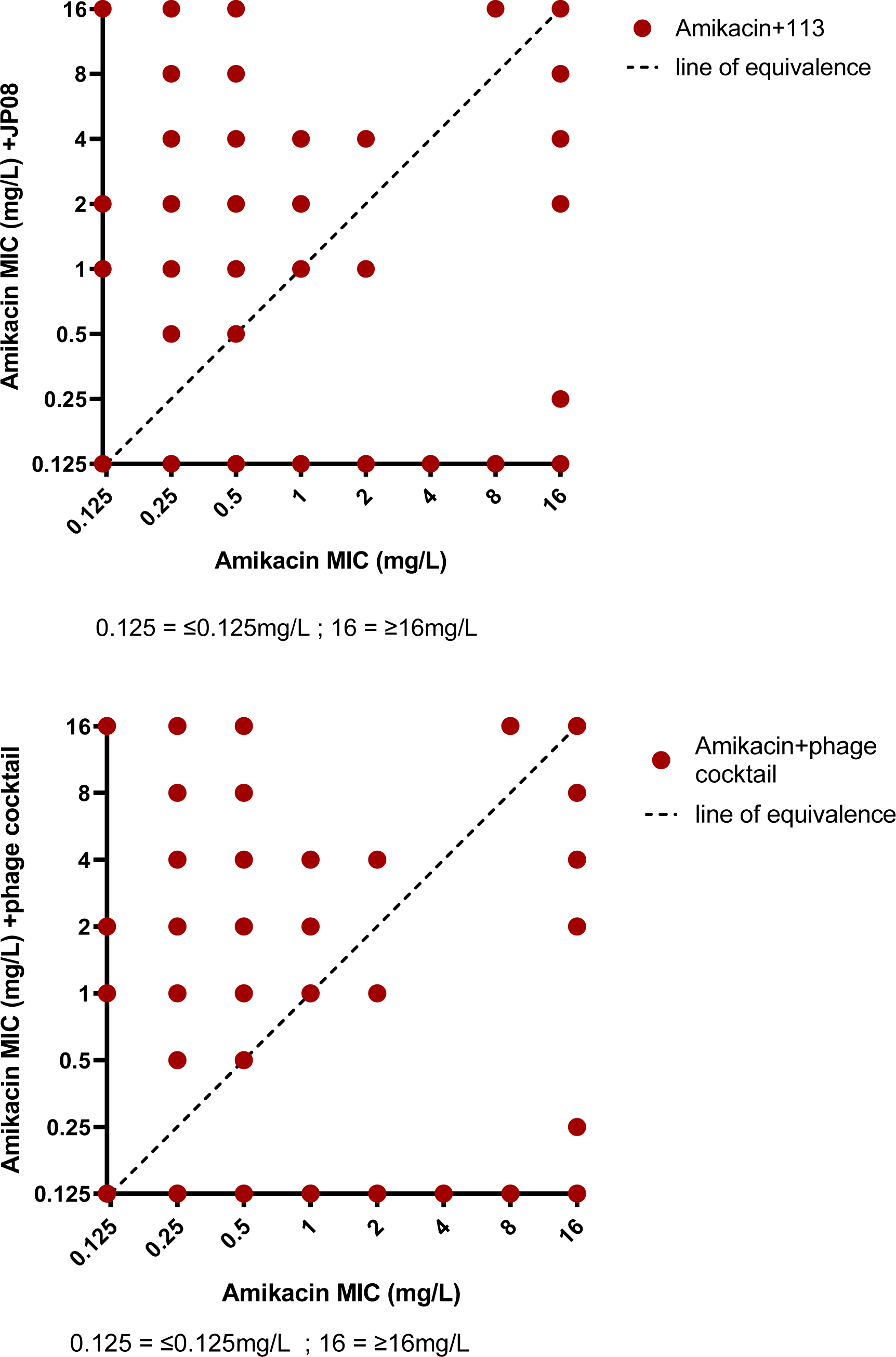
Plots of amikacin MIC versus amikacin MIC in the presence of phage.

**Figure S2).**
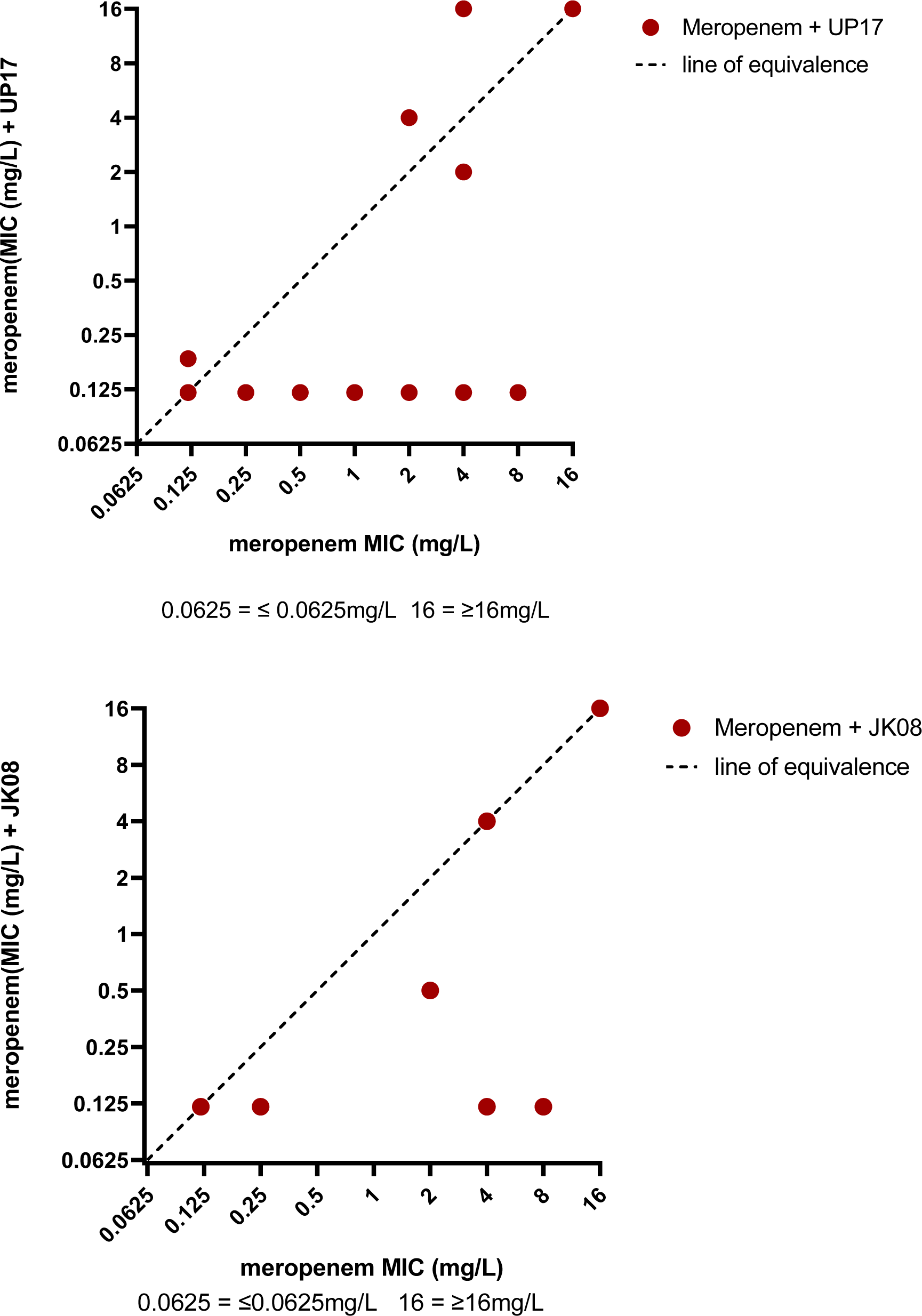
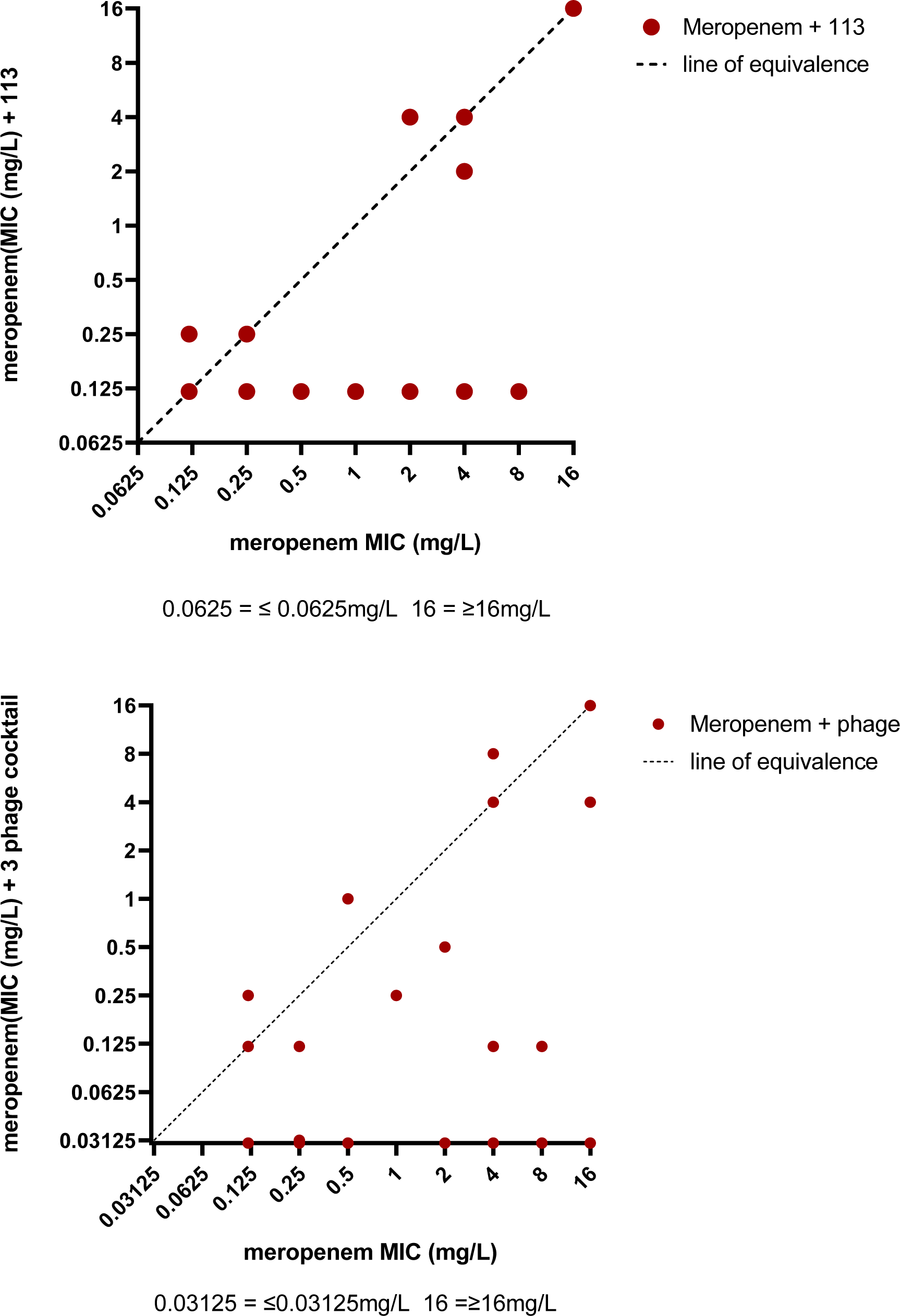
Plots of meropenem MIC versus amikacin MIC in the presence of phage.

## References

1) Palmer JD, Foster KR. The evolution of spectrum in antibiotics and bacteriocins. Proc. Natl. Acad. Sci. U.S.A 2022;119, e2205407119.

2) Amabile-Cuevas CF. Phage therapies: Lessons (Not) learned from the ‘Antibiotic Era’. Phage 2022;3, 10.1089/phage.2022.0001

3) World Health Organsation Antimicrobial resistance division, National action plans and monitoring and Evaluation. Global Action Plan on Antimicrobial Resistance 2016 ISBN 9789241509763 https://who.int/publications/i/item/9789241509763

4) Chaudhry WN, Concepción-Acevedo J, Park T, et al. Synergy and order effects of and in killing *Pseudomonas aeruginosa* biofilms. PLoS One. 2017;12:e0168615.

5) Pirnay J-P, Djebara S, Steurs G et al Retrospective observational analysis of the first one hundred consecutive cases of personalized bacteriophage therapy of difficult to treat infections facilitated by a Belgian consortium. medRxiv 2023.08.28.23294728; 10.1101/2023.08.28.23294728

6) International Organistion for Standardization 2019 Susceptibility testing of infectious agents and evaluation of performance of antimicrobial testing devices - Part 1 http://www.iso.org/obp/ul/#iso:std:15020776-1:ed-2:v2:en

7) EUCAST Clinical Breakpoint Table v14.0 accessed 08/01/2024 https://www.eucast.org/clinical_breakpoints

8) Noel AR, Attwood M, Bowker KE, et al Comparative bactericidal activity of representative β-lactams against Enterobacterales, Acinetobacter baumannii and Pseudomonas aeruginosa, J Antimicrobial Chemother, 2022; 77, 1306–1312, 10.1093/jac/dkac026

9) Liu CG, Green SI, Min L et al Phage-Antibiotic Synergy Is Driven by a Unique Combination of Antibacterial Mechanism of Action and Stoichiometry. mBio 2020:11 11:e01462–20. 10.1128/mBio.01462-20.

10) Attwood M, Griffin P, Noel AR, et al Antibacterial effect of seven days exposure to ceftolozane-tazobactam as monotherapy and in combination with fosfomycin or tobramycin against Pseudomonas aeruginosa with ceftolozane-tazobactam MICs at or above 4 mg/l in an in vitro pharmacokinetic model. J Antimicrob Chemother. 2023 ;78:2254–2262. 10.1093/jac/dkad230.

11) Dufour N, Delattre R, Ricard J-D et al The Lysis of Pathogenic Escherichia coli by Bacteriophages Releases Less Endotoxin Than by β-Lactams. Clin Infect Dis. 2017;64:1582–1588. doi: 10.1093/cid/cix184 Erratum in: Clin Infect Dis. 2017;65(8):1431–1433.

12) Knezevic P, Curcin S, Aleksic V et al Phage-antibiotic synergism: a possible approach to combatting Pseudomonas aeruginosa Research in Microbiology, 2013; 164:55–60. 10.1016/j.resmic.2012.08.008.

13) MacGowan A, Rogers C, Bowker K In Vitro Models, In Vivo Models, and Pharmacokinetics: What Can We Learn from In Vitro Models?, Clinical Infectious Diseases 2001; 33 Supplement 3, S214–S220, 10.1086/321850

14) Dąbrowska K, Abedon ST. Pharmacologically Aware Phage Therapy: Pharmacodynamic and Pharmacokinetic Obstacles to Phage Antibacterial Action in Animal and Human Bodies. Microbiol Mol Biol Rev. 2019;83:e00012–19. 10.1128/MMBR.00012-19.

